# Reduced RNA turnover as a driver of cellular senescence

**DOI:** 10.1101/800128

**Authors:** Nowsheen Mullani, Yevheniia Porozhan, Mickael Costallat, Eric Batsché, Michele Goodhardt, Giovanni Cenci, Carl Mann, Christian Muchardt

## Abstract

Accumulation of senescent cells is an important contributor to chronic inflammation upon aging. While cytoplasmic DNA was shown to drive the inflammatory phenotype of senescent cells, an equivalent role for RNA has never been explored. Here, we show that some senescent cells accumulate long promoter RNAs and 3’ gene extensions, rich in retrotransposon sequences. Accordingly, these cells display increased expression of genes involved in detecting double stranded RNA of viral origin downstream of the interferon pathway. The RNA accumulation is correlated with signs of reduced RNA turn-over, including in some cases, reduced expression of RNA exosome subunits. Reciprocally, engineered inactivation of RNA exosome subunit Exosc3 induces expression of multiple senescence markers. A senescence-like RNA accumulation is also observed in cells exposed to oxidative stress, an important trigger of cellular senescence. Altogether, we propose that in a subset of senescent cells, repeat-containing transcripts stabilized by oxidative stress or reduced RNA exosome activity participate, possibly in combination with cytoplasmic DNA, in driving and maintaining the permanent inflammatory state characterizing cellular senescence.

## INTRODUCTION

Cellular senescence is a state of irreversible cell cycle arrest (Rodier & Campisi, 2011). Experimentally, it can be triggered via multiple routes including prolonged maintenance in tissue culture, exposure to ionizing radiations or oxidative stress, and forced expression of mitogenic oncogenes. These inducers of senescence all share the ability to cause DNA damage and to generate reactive oxygen species, two components that probably are at the basis of the phenomenon (Ben-Porath & Weinberg, 2005). One of the hallmarks of senescent cells is their production of a range of chemokines, proinflammatory cytokines, growth factors, and matrix-remodelling enzymes, defining the senescence-associated secretory phenotype or SASP. This proinflammatory characteristic has a crucial role in propagating senescence and in recruiting immune cells to the senescent tissue. As senescent cells accumulate with time, the SASP is also believed to be a major determinant of the chronic low-grade inflammation associated with ageing and age-related diseases.

SASP activation is largely orchestrated by nuclear factor NF-kB and CCAAT/enhancer-binding protein beta (C/EBPb). Upstream of these transcription factors, DNA damage and the DNA damage response (DDR) are majors triggers of the proinflammatory pathways (Salminen, Kauppinen et al., 2012). Yet, it seems that other mechanisms may allow nucleic acids to drive the chronic sterile inflammation characteristic of cellular senescence. Indeed, several studies have associated senescence with an accumulation of DNA in the cytoplasm. This DNA, in the form of chromatin, triggers innate immunity via the cytosolic DNA-sensing cGAS–STING pathway (Dou, Ghosh et al., 2017). Cytoplasmic DNA possibly originates from chromosome segregation errors during mitosis and its accumulation seems favoured by down-regulation in senescent cells of the cytoplasmic DNases TREX1 and DNASE2 (Takahashi, Loo et al., 2018). In addition, it has been shown that in senescent cells, de-repression of repeat elements of the LINE family results in the production of retroviral RNAs, that after retrotranscription, accumulate in the cytoplasm in the form of cDNAs (Cecco, Ito et al., 2019).

Consistent with a role of cytoplasmic DNA, multiple studies have documented the importance of the interferon pathway in driving senescence, and suppression of type 1 interferon signaling hinders the onset of senescence (Katlinskaya, Katlinski et al., 2016). The interferon pathway was initially described as an antiviral defense mechanism activated by specific cytoplasmic or endosomal receptors of either viral DNA or double stranded RNA. In this context, RNA could also be a trigger of senescence. However to date, the role of RNA in senescence has mostly been examined in the nuclear compartment, where specific long non-coding RNAs (lncRNAs) participate in the regulation of genes relevant for cellular senescence (Du, Yang et al., 2016a, Du, Yang et al., 2016b, Lazorthes, Vallot et al., 2015). Nevertheless, some observations are suggestive of a wider impact of RNAs in cellular senescence, via their production, their maturation, or their turn-over. For example, it was reported that neurons in the aging mouse brain accumulate 3′ untranslated regions (UTRs), resulting in the production of small peptides of yet unknown function (Sudmant, Lee et al., 2018). Furthermore, transcripts from SINEs/Alus, a family of repetitive elements particularly abundant in euchromatin, were reported to accumulate in senescent cells with an impact on genome integrity (Wang, Geesman et al., 2014). Finally, several studies have reported that cellular senescence is associated with numerous changes in the outcome of alternative pre-mRNA splicing (Wang, Wu et al., 2018). This results in the production of senescent toxins including progerin, a variant of Lamin A associated with the Hutchinson-Gilford progeria syndrome (Cao, Blair et al., 2011), while also favoring the synthesis of S-Endoglin and p53beta that may have similar pro-senescence activities (Fujita, Mondal et al., 2009, Miao, Wu et al., 2016).

Whether RNAs have an effect on cellular senescence independently of their ability to regulate specific genes in the nucleus or to encode proteins has never been specifically investigated. Here, we examined RNA-seq data from senescent cells of different origins for sources of RNA species liable to activate innate immunity pathways. This approach revealed that in several cell lines, senescence is associated with a gradual accumulation of reads originating from regions located upstream or downstream of genes and extending outside of the transposon-free regions characterizing transcription start and termination sites. Accumulation of these RNAs that are all substrates of the RNA exosome, was correlated with indications of reduced RNA turnover and increased activity of anti-viral pathways dedicated to the detection of double-stranded RNAs. Consistent with a pro-senescence activity of the accumulating RNAs, we further showed that inactivation of RNA exosome activity in mouse cells gives rise to senescence-like characteristics. As oxidative stress causes an accumulation of RNA species largely resembling that observed in senescent cells, we propose that reactive oxygen species of mitochondrial origin may be the trigger of an RNA-dependent mechanism possibly involved in the permanent inflammatory state of senescent cells.

## RESULTS

### Accumulation of RNA exosome substrates in senescent Wi38 cells

It was previously reported that in Wi38 cells driven into senescence by oncogenic RAF, many convergent genes displayed transcriptional termination read-through resulting in transcriptional interference (Muniz, Deb et al., 2017). In the RNA-seq data from this study, the read-out of the read-through was an accumulation of sequencing reads downstream of genes in the senescent cells. Re-analysis of the data revealed that this accumulation occured at the wide range of genes beyond those described in the initial study. In addition, examination of canonical histone genes, transcribed mainly in the S phase of the cell cycle and therefore strongly downregulated in senescent cells, suggested that accumulation of downstream reads was not correlated with the activity of the cognate gene (Figure 1A, green arrows).

**Figure 1:**
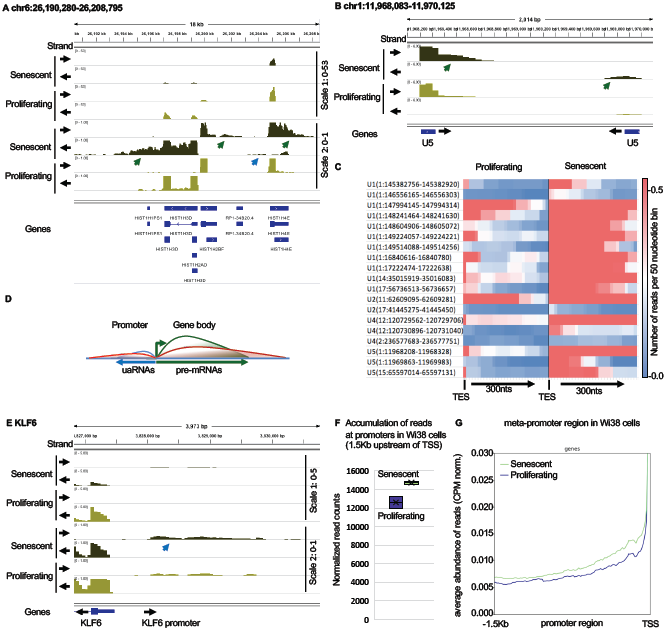
Senescent WI38 cells accumulate pegeRNAs. RNA-seq data from WI38 hTERT RAF1-ER human fibroblasts either proliferating or driven into senescence by induction of RAF1-ER (Lazorthes et al., 2015). (A, B, E) RNA-seq reads were quantified, normalized by the CPM method and displayed with IGV. Black arrows indicate the orientation of the transcription. Green arrows indicate 3’ extensions, blue arrows, uaRNAs. (C) Heatmap illustrating increased accumulation of 3’ extensions of U snRNAs in senescent vs. proliferating WI38 hTERT RAF1-ER cells. Counts are normalized by the CPM method. TES indicates the 3’ end of the U snRNA gene. The U snRNAs are also listed in Sup. Table 1A. (D) Schematic representation of divergent transcription at promoters. Green line represents pre-mRNA, blue line, normal accumulation of uaRNAs, red line, accumulation of uaRNAs in senescent cells. (F) At 5260 human promoters not overlapping with coding regions of any gene, reads were counted (CPM normalization) within a region of 1500 nucleotides upstream of the transcription start site (TSS) in either proliferating or senescent WI38 hTERT RAF1-ER cells. (G) Average profile of read distribution along the 5260 promoters in proliferating and senescent WI38 hTERT RAF1-ER cells (CPM normalization).

RNAs transcribed downstream of genes are normally degraded by the RNA exosome, upon cleavage of the main transcript by the cleavage and polyadenylation apparatus (CPA) for most genes, or by a dedicated machinery in the case of the canonical histone genes. Consistent with a role for the RNA exosome in the degradation of 3’ extensions of histone genes, we noted that depletion of EXOSC3 in HeLa cells resulted in an accumulation of downstream reads very similar to that observed in the senescent Wi38 cells (Sup. Figure 1A, green arrows).

The similarity in the profile of downstream reads in cells depleted for EXOSC3 and senescent cells prompted us to explore whether senescent cells accumulate other RNA exosome substrates. Interestingly, we found increased levels of 3’ extensions at essentially all U snRNA genes expressed in the Wi38 cells (Figure 1B and 1C). These 3’ extensions are normally cleaved by the Integrator complex (Baillat, Hakimi et al., 2005) and accumulate when the RNA exosome is inactivated (Sup. Figure 1B). Re-observation of the histone genes further revealed an accumulation of upstream antisense RNAs (uaRNAs) at their promoters (blue arrow, Figure 1A). These uaRNAs are produced as a consequence of the divergent transcription occurring at sites of transcription initiation (schematic Figure 1D). Consistent with a degradation of uaRNAs by the RNA exosome, inactivation of EXOSC3 in HeLa cells resulted in a similar accumulation of uaRNAs at the histone genes (blue arrow Sup. Figure 1A and Sup. 1C). Further examination of the Wi38 RNA-seq data showed that senescence correlated with accumulation of uaRNAs at many additional genes (see example of the KLF6 gene Figure 1E and 119 additional examples in Sup. Table 1A). Quantification at a series of 5260 promoters not overlapping with coding regions of any gene revealed an approximately 10% increase in uaRNA accumulation in the senescent cells (Fig 1F and 1G).

Together, these observations are highly suggestive of a reduced activity of the RNA exosome of Wi38 oncogene-induced senescence, resulting in the accumulation of transcripts originating from regions upstream and downstream of genes. We will refer to these RNAs as perigenic RNAs (pegeRNAs).

### Reduced expression of RNA exosome catalytic subunit in senescent Wi38 cells

To identify possible causes for the accumulation of RNA exosome substrates in the senescent Wi38 cells, we compared their transcriptomes to that of proliferating Wi38 cells, with a particular interest in the expression of RNA exosome subunits. In these senescent cells, we observed a strongly decreased expression of DIS3L, the catalytic subunit of the cytoplasmic RNA exosome complex (Figure 2A). This decreased expression resulted in decreased accumulation of the DIS3L protein, while levels of the senescence marker CDKN1A/p21 were increased (Figure 2B). Of note, the regulatory RNA exosome subunit EXOSC4 was upregulated in these cells (Figure 2A).

**Figure 2:**
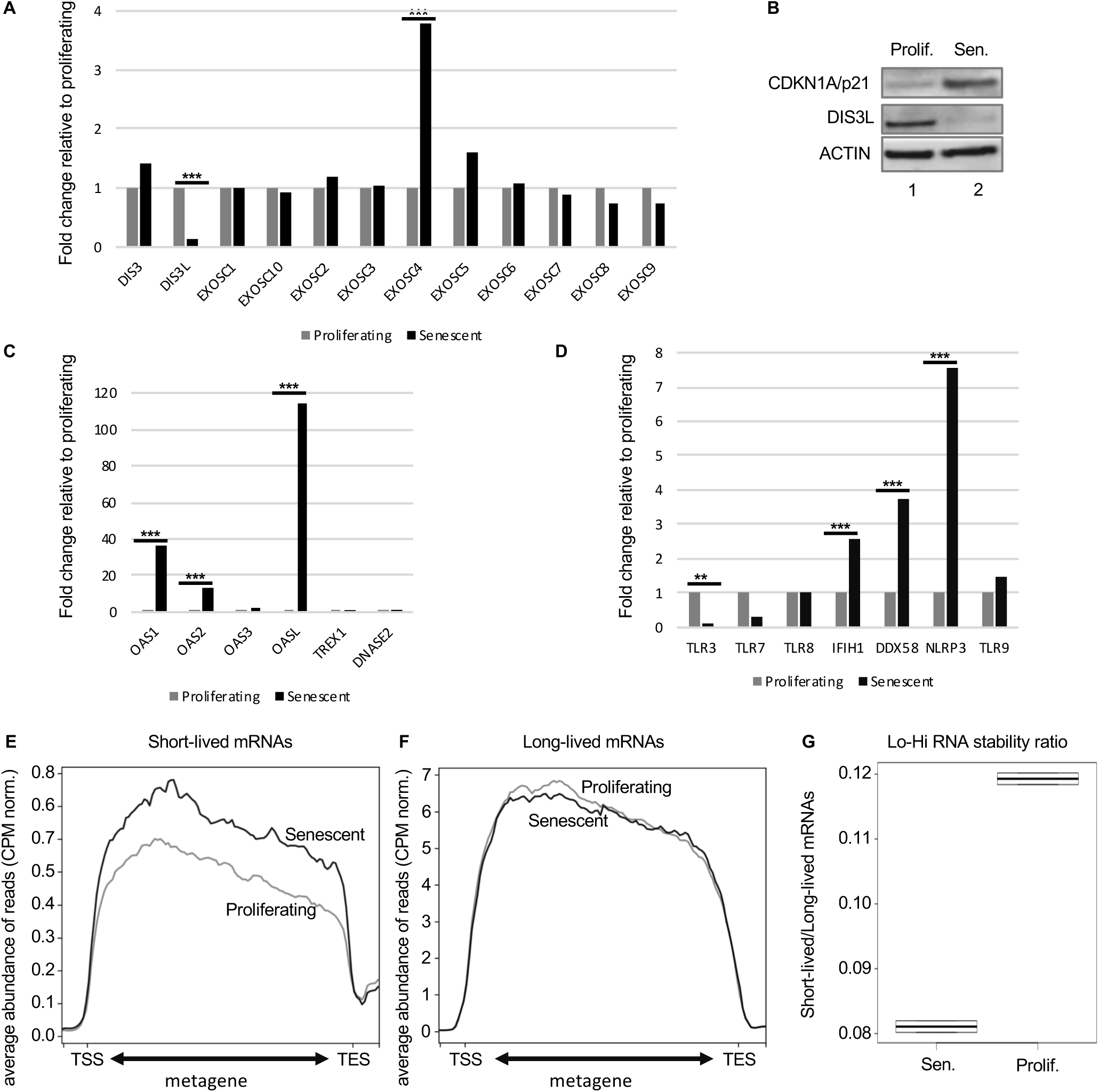
Reduced expression of RNA exosome subunits in senescence. (A, C, D) Differential gene expression was estimated with DESeq2 on RNA-seq data (n=2) from WI38 hTERT RAF1-ER human fibroblasts either proliferating or driven into senescence by induction of RAF1-ER (Lazorthes et al., 2015). Histograms represent fold change in expression relative to proliferating cells for the indicated genes. *** and ** indicate p-values below 0.001 and 0.01 respectively as calculated by DESeq2. (B) Western blots with indicated antibodies on total extracts from either proliferating (lane 1) or driven into senescence by induction of RAF1-ER (lane 2). (E, F) Average profile of reads mapping to short-lived (less than 2h) or long-lived (more than 10h) mRNAs as listed in (Tani et al., 2012), either in proliferating (light grey) or in senescent (dark grey) cells as indicated. Read counts were normalized by CPM. (G) Ratio of the number of reads mapping to short-lived (less than 2h) over the number of reads mapping to long-lived (more than 10h) mRNAs.

We then examined the transcriptomes for signs of activation of defense mechanisms against cytoplasmic RNA, first with focus on the anti-viral interferon-activated OAS/RNASEL pathway. In this pathway, the 2′,5′-oligoadenylate synthetases OAS1, OAS2, and OAS3 generate a 2-5A modification on double-stranded RNAs, making them targets for RNASEL-mediated degradation (Silverman, 2007). Consistent with an activation of this pathway in the senescent cells, we observed an increased expression of both OAS1 and OAS2 (Figure 2C). The OASL gene, encoding a catalytically inactive isoform that retains double stranded RNA binding activity, displayed an even stronger activation. In contrast, TREX1 and DNASE2, the two DNAses previously described as having a reduced expression in some senescent cells (Takahashi et al., 2018) were unaffected here. Finally, we examined variations in expression of cytoplasmic or endosomal receptors for RNA (including TLR3, TLR7, TLR8, IFIH1/MDA5, DDX58/RIG-I, and NLRP3), and for DNA (TLR9). Among these, IFIH1/MDA5, DDX58/RIG-I and particularly NLRP3, the sensor component of the NLRP3 inflammosome, were significantly upregulated, appearing as potential candidates for the detection of pegeRNAs (Figure 2D).

To gain further evidence for a reduced RNA turnover in the senescent Wi38 cells, we compared accumulation of short-lived and long-lived mRNAs, the assumption being that unstable RNAs would be more affected by fluctuation in RNA exosome activity than would the stable mRNAs. For this, we established a list of highly unstable (t_1/2_<2h) or highly stable (t_1/2_>10h) mRNAs from a genome-wide study on mRNA stability (Tani, Mizutani et al., 2012). We first verified our assumption by examining the effect of reduced RNA exosome activity on accumulation of these mRNAs in HeLa cells. This showed that depletion of EXOSC3 resulted in increased accumulation of the short-lived RNAs when compared to WT cells. In contrast, levels of long-lived mRNAs appeared moderately decreased (Sup. Figure 2A-2B). We explain this decrease by the augmented complexity of the RNA samples in the absence of RNA exosome activity, that may result in less RNA-seq reads per RNA species. The final outcome of these variations was a 30-40% increase (pVal<0.05) in the ratio of the number of reads aligning on short-lived mRNAs over those aligning on long-lived mRNAs (Sup. Figure 2C). Furtheron, we will refer to this ratio as the Lo-Hi RNA stability ratio. When applied to the Wi38 data, this approach revealed that the short-lived mRNAs accumulated more in senescent cells than in the proliferating cells, while levels of long-lived mRNAs remained unchanged between the two conditions (Figure 2E-2F). The resulting Lo-Hi RNA stability ratio was increased by approximately 40%, similar to that observed in cells depleted for EXOSC3 (Figure 2G and Sup. Figure 2C).

Together, these observations argued in favor of a reduced turnover of unstable RNAs in the senescent Wi38 cells, associated with increased activity of innate immune defense mechanisms implicated in defense against RNA viruses.

### Reduced turn-over of unstable RNAs is observed in multiple senescent cells

To investigate whether the reduced turn-over of unstable RNAs observed in the Wi38 cells driven into senescence by oncogenic RAF was an exception or a widespread phenomenon, we examined several additional RNA-seq data sets from senescent cells.

First, we examined another case of oncogene induced senescence involving expression of oncogenic RAS in IMR90 human fibroblasts. In this series, RNA-seq (n=2) was carried out after expression of RAS for 0, 4, or 10 days (Lau, Porciuncula et al., 2019). Examination of the transcriptome of the cells revealed a progressive reduction in transcripts from several genes encoding subunits of the RNA exosome, including DIS3L, EXOSC2, EXOSC3, EXOSC6, EXOSC8, EXOSC9, and EXOSC10 (Figure 3A). As in the Wi38 cells, we also noted an upregulation of the regulatory subunit EXOSC4. This data set was of insufficient depth to visualize pegeRNAs, we therefore examined (1) the accumulation of unstable compared to stable mRNAs, and (2) the expression of the OAS enzymes involved in detecting and degrading nucleic acids in the cytoplasm, as described above. Consistent with a reduced RNA turnover, we found that the Lo-Hi RNA stability ratio increased progressively during the entry of the IMR90 cells into senescence (Figure 3B-3C). OAS1 and OAS2 mRNA levels increased accordingly. The DNAse TREX1 was moderately downregulated, while DNASE2 was unexpectedly upregulated, as was the DNA receptor TLR9. The possible interpretation of the variations of DNASE2 and TLR9 expression will be addressed in the discussion.

**Figure 3:**
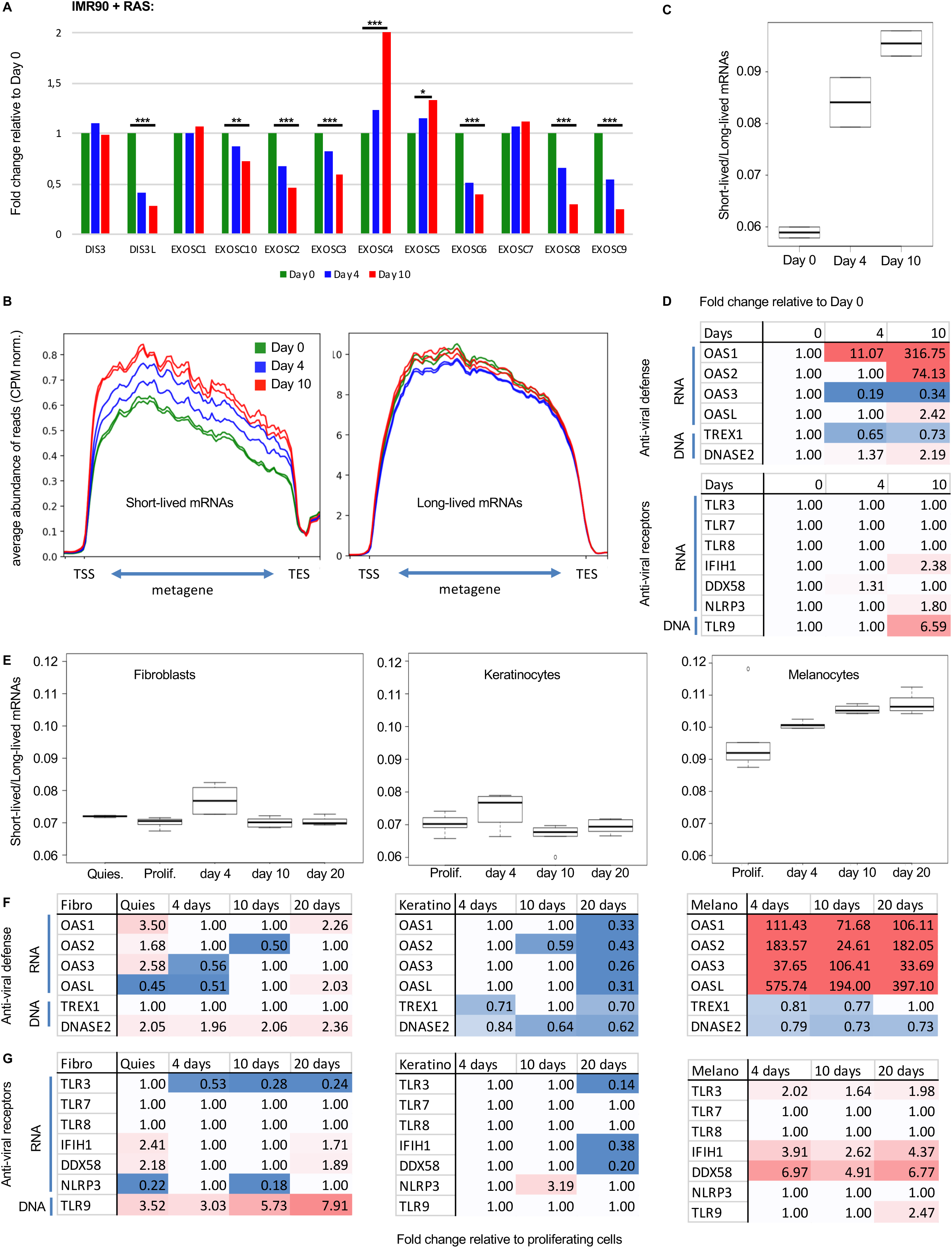
Reduced RNA decay in senescent cells of various origin. (A-D) RNA-seq data from IMR90 human fibroblasts induced to senescence via activation of the oncogene Ras (Lau et al., 2019). Samples (n=2) had been collected at day 0 (growing phase), day 4 (beginning of SASP induction) and day 10 (senescent phase) of Ras induction. (A) Differential gene expression was estimated with DESeq2. Histograms represent fold change in expression relative to proliferating cells for the indicated genes. *** and ** indicate p-values below 0.001 and 0.01 respectively as calculated by DESeq2. (B) Average profile of reads (CPM normalization) mapping to short-lived (less than 2h) or long-lived (more than 10h) mRNAs as in Figure 2 at the indicated time points. (C) Lo-Hi RNA stability ratio for the experiment at the indicated time points, calculated as in Figure 2. (D) Fold change relative to day 0 of the indicated genes at the indicated time point. Red gradient indicates upregulation (max red at 10-fold), blue indicates downregulation (max blue at 0.5). Variations with pVal>0.05 were set to 1. (E-F) RNA-seq data from (Hernandez-Segura et al., 2017), n=6 for each cell type and time point. HCA-2 (fibroblasts), keratinocytes, or melanocytes had been exposed to ionizing radiation. RNA had been harvested from proliferating cells, from cells brought into quiescence (Quies) by culture for 48h in the presence of 0.2%FBS, or 4, 10, or 20 days after irradiation. (E) Lo-Hi RNA stability ratio for each experiment calculated as indicated in Figure 2. (F-G) Fold change relative to proliferating cells of the indicated genes calculated and represented as in (D).

Next, we examined a dataset from human fibroblasts (HCA-2 cells), keratinocytes, and melanocytes driven into senescence by ionizing radiation (n=6, with data points at 0, 4, 10, and 20 days of culture) (Hernandez-Segura, de Jong et al., 2017). In this series, the Lo-Hi RNA stability ratio was gradually increased only in the melanocytes (Figure 3E). These cells were also the only ones to upregulate the OAS genes, consistent with them having to cope with accumulation of double stranded RNAs (Figure 3F). Transcription of the RNA receptors TLR3, IFIH1/MDA5, and DDX58/RIG-I was also increased at all timepoints in the melanocytes, while the fibroblasts upregulated the DNA receptor TLR9 (Figure 3G). Of note, in the melanocytes, it was not possible to unambiguously associate the suspected reduced RNA turnover with reduced transcription of any specific RNA exosome gene. This will be further addressed in the discussion. Finally, this data set also included RNA-seq from quiescent fibroblasts, allowing us to verify that reduced RNA turnover was not triggered by simple growth arrest (Figure 3E, 3F, 3G, Quies).

To further probe the robustness of our observations, we finally examined a large dataset from Wi38, IMR90, HAEC, and HUVEC cells driven into senescence via different routes (n=2 for each condition), including oncogenic RAS, replicative exhaustion, and DNA damage by treatment with either doxorubicin or gamma irradiation (Casella, Munk et al., 2019). In all experiments on Wi38 cells, senescence led to an increased Lo-Hi RNA stability ratio (Sup. Figure 3A). Consistent with this, expression the OAS genes was upregulated upon replicative and DNA damage-induced senescence (Sup. Figure 3B, top panel). This activation of the OAS genes correlated with increased expression of the TLR3, IFIH1/MDA5, and DDX58/RIG-I sensors of double stranded RNA (Sup. Figure 3B, bottom panel). Interestingly, in the RAS-induced Wi38 senescence, like in the initial RAF-induced Wi38 senescence (Figure 2D), strongest transcriptional activation was observed for the catalytically inactive OASL and the inflammosome component NLRP3 (Sup. Figure 3B, column 1). In all cases, the DNases TREX1 and DNASE2 were affected less than 2-fold although we noted a 3-4-fold activation of the DNA receptor TLR9 in two of the data sets (Dox(2) and 10Gy).

From the same dataset, we next analysed the RNA-seq from 3 different cells types, HAEC, HUVEC and IMR90 cells driven into senescence by gamma irradiation or replicative exhaustion. We observed that the Lo-Hi RNA stability ratio was only moderately increased for the HAEC and HUVEC cells, while it was decreased in the IMR90 cells. The modestly increased Lo-Hi RNA stability ratio in the HAEC and HUVEC cells translated into a small but significant (pVal<0.05) increase in OAS1, OAS2, TLR3, and IFIH1/MDA5 gene expression in the HAEC and HUVEC cells, while these genes were unaffected or down-regulated in the IMR90 cells.

Altogether, observations of these cellular models indicate that it is possible to define a subcategory of senescent cells displaying increased accumulation of the more unstable RNAs and signs of a defense reaction against double stranded RNA. We speculate that in this subcategory, reduced turnover and subsequent accumulation of specific RNA species participate, possibly together with cytoplasmic DNA, in triggering the inflammatory phenotype associated with senescence.

### Senescent cells share an RNA signature with cells exposed to oxidative stress

Accumulation of uaRNAs has previously been observed in cells exposed to oxidative stress, a central determinant of senescence ((Giannakakis, Zhang et al., 2015, Nilson, Lawson et al., 2017) and example of uaRNAs at the KLF6 locus, Sup. Figure 4A and 4B). This phenomenon was attributed to defective RNA polymerase II transcription termination (Nilson et al., 2017). To investigate possible similarities between oxidative stress and senescence at the level of the RNAs accumulating in the cells, we examined cells exposed to H2O2 for the presence of pegeRNAs other than uaRNAs. For this study, we chose a data set from H2O2-treated BJ or MRC5 cells allowing for detection of rare RNAs (Giannakakis et al., 2015). In this data, like in the RAF-induced Wi38 senescent cells and the EXOSC3-depleted HeLa cells, we observed an accumulation of reads downstream of many histone genes (Figure 4A and Sup. Figure 4C). We also observed the accumulation of 3’ extensions of U snRNAs (see example in Figure 4B and heatmap of Figure 4C). Finally, as in the senescent cells, we observed a gradual increase in the Lo-Hi RNA stability ratio, suggestive of a reduced RNA turnover (Figure 4D and Sup. Figure 4E).

**Figure 4:**
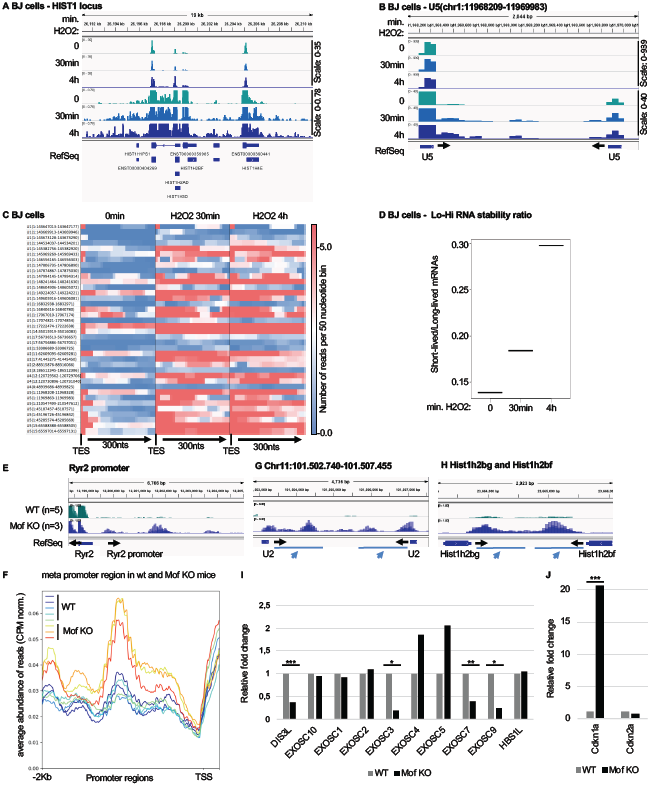
Accumulation of pegeRNAs in cells exposed to oxidative stress. (A-D) BJ human fibroblasts exposed to 0.2mM H2O2 for the indicated times (one sample per time point) (Giannakakis et al., 2015). (A, B) RNA-seq reads were quantified, normalized by the CPM method and displayed with IGV. (C) Heatmap illustrating increased accumulation (CPM normalization) of non-maturated U snRNAs in BJ cells exposed to H2O2 for the indicated times. TES indicates the 3’ end of the U snRNA gene. The U snRNAs are also listed in Sup. Table 2. (D) Lo-Hi RNA stability ratio for BJ cells at the indicated time points, calculated as indicated in Figure 2. (E-J) Mouse cardiomyocytes either WT (n=5) or inactivated for the Mof histone acetylase (n=3) were analyzed by RNA-seq on poly(A)-enriched libraries (Chatterjee et al., 2016). (E, G, H) RNA-seq reads were quantified, normalized by the CPM method and displayed with IGV genome browser. Overlay of several track as indicated. (F) Average profile of read distribution (CPM normalized) along the promoters of 1200 mouse genes with similar expression levels in either WT or Mof KO cardiomyocytes. (I, J) Differential gene expression was estimated with DESeq2. Histograms show variations of the indicated genes. ***, **, and * indicate p-values below 0.001, 0.01, and 0.05 respectively.

We next examined the transcriptome of a mouse model involving oxidative stress associated with mitochondrial dysfunction caused by inactivation of the Mof histone acetylase in the heart (Chatterjee, Seyfferth et al., 2016). Inactivation of this gene has catastrophic consequences on cardiac tissue that has high-energy consumption, triggering hypertrophic cardiomyopathy and cardiac failure. At a cellular level, cardiomyocytes were reported to show severe mitochondrial degeneration and deregulation of mitochondrial nutrient metabolism and oxidative phosphorylation pathways. Consistent with mitochondrial suffering and ensuing oxidative stress, Mof inactivation reproduced the increased accumulation of uaRNAs observed in the H2O2-treated human tissue-culture cells (see example of the Ryr2 promoter, Figure 4E, and metaplot of 1200 promoters, Figure 4F). Likewise, Mof inactivation resulted in accumulation of 3’ extensions at U snRNA and histone genes, reproducing another RNA signature of senescence (Figure 4G and 4H, note that the RNA-seq reactions were based on poly(A)-enriched libraries, preventing detection of mature U snRNAs and histone mRNAs). Consistent with reduced RNA decay, we noted in the Mof KO cells, a significant decrease in the expression of several subunits of the RNA exosome, particularly the catalytic subunit Dis3l, as well as Exosc3, Exosc7, and Exosc9 (Figure 4I). In parallel, and consistent with Mof inactivation causing oxidative stress and DNA damage, we noted a clear increase in the expression of the senescence marker Cdkn1a/p21 (Figure 4J).

Together, these observations suggest that the oxidative stress associated with cells undergoing senescence may be upstream of the accumulation of pegeRNAs observed in these cells.

### RNA exosome depletion induces characteristics of senescence in mouse ES cells

With the objective of identifying a possible causative role of pegeRNAs in the onset of senescence, we next explored the biological effect of their accumulation in primary cells. To this end, we examined a dataset from mouse embryonic stem cells depleted in Exosc3 for 3 days (Chiu, Suzuki et al., 2018). As expected and as described in the original paper, these cells recapitulated the increased accumulation uaRNAs (example in Figure 5A). They also recapitulated the effect on U snRNA maturation (example in Figure 5B). Interestingly, GO term analysis of genes up-regulated upon Exosc3 inactivation revealed a highly significant enrichment in genes associated with the p53 pathway and we pinpointed an increased expression of both Cdkn1a (p21) and Cdkn2a (p16), two markers of cellular senescence (Figure 5C and 5D, and Sup. Table 5 listing differentially regulated genes associated with the GO terms). Consistent with this, downregulated genes were highly enriched in genes associated with cell cycling (Figure 5E). From these observations, we concluded that loss of RNA exosome activity resulted in a transcriptional landscape comparable to that of senescent cells.

**Figure 5:**
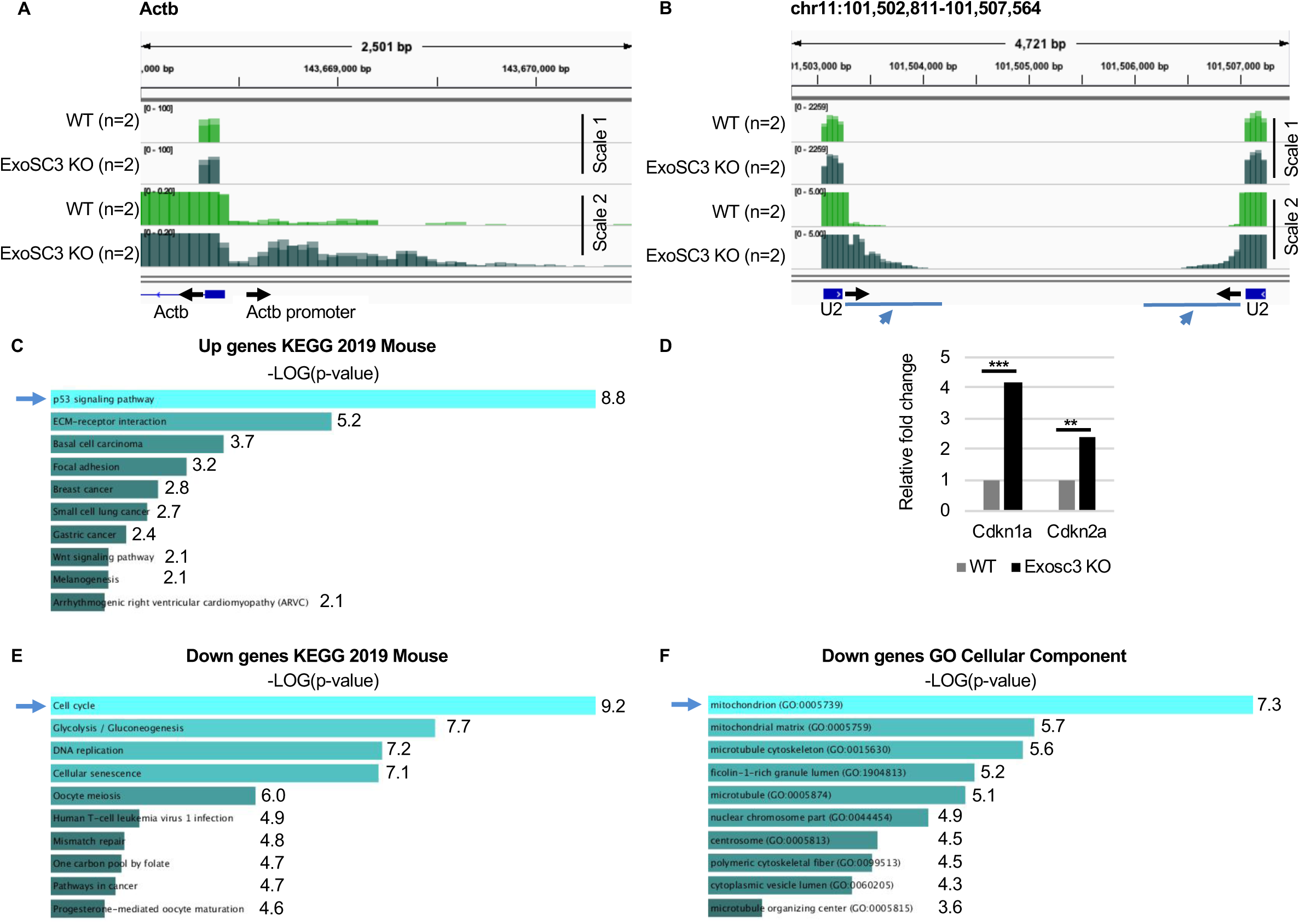
Inactivation of the RNA exosome induces senescence-like features and deregulates mitochondrial genes. RNA-seq data (n=2) from mouse ES cells inactivated for Exosc3 and harboring an inducible Exosc3 expression construct human (Chiu et al., 2018). Exosc3 expression is initially induced (WT), then the inducer is removed from the medium and cells are cultured for 3 days (Exosc3 KO). (A, B) RNA-seq reads were quantified, normalized by the CPM method and displayed with IGV genome browser. Black arrows indicate the orientation of the gene. Blue arrows indicate regions of interest. Overlay of several track as indicated. (C-F) Differential gene expression was estimated with DESeq2. (C, E, F) GO term analysis of genes significantly affected by Exosc3 inactivation (p-values<0.05) was carried out with Enrichr. Histograms represent -LOG(p-value) of the enrichment in each indicated path. (D) Histograms represent fold change in expression relative to WT cells for the indicated genes. *** and ** indicate p-values below 0.001 and 0.01 respectively as calculated by DESeq2.

Examining GO terms for cellular compartments further highlighted a significant enrichment in mitochondrial genes among the genes down-regulated by depletion of Exosc3 (Figure 5F). This was consistent with an earlier report showing mitochondrial dysfunction in patients with pontocerebellar hypoplasia, a syndrome linked to a mutation in the EXOSC3 gene (Schottmann, Picker-Minh et al., 2017).

Together, these observations indicate that reduced RNA exosome activity has physiological consequences that resemble cellular senescence. In addition, the data are suggestive of a bidirectional crosstalk between RNA degradation and oxidative stress for the induction of growth arrest.

### Serendipitous transcription of DNA repeats in pegeRNAs

We finally investigated why pegeRNAs would have the potential to trigger cytoplasmic anti-viral mechanisms. In that context, we noted that an earlier study had reported an accumulation in senescent cells of RNAs originating from the SINE family of retrotransposons (Wang et al., 2014). In parallel, another study showed that Alu-containing RNAs can trigger an interferon response and stimulate secretion of cytokines (Hung, Pratt et al., 2015). Together, these observations prompted us to envision that pegeRNAs may be enriched in SINE and possibly other repeated sequences prone to fold into double-stranded RNAs with triggering effects on the interferon pathway.

To examine this possibility, we focused on LINEs and SINEs (including Alus) that are the most abundant repeated sequences in the vicinity of genes. Plotting the profile of distribution of sequences annotated “SINE” or “LINE” in RepeatMasker relative to protein-coding genes (annotated NM in RefSeq) showed that an average gene is framed by two short repeat-depleted regions before the transcription start (TSS) and after the transcription end (TES) sites (Fig. 6A). Likewise, the profile of SINE and LINE distribution over U snRNA genes showed that only the gene bodies are depleted of the repeated sequences (Fig. 6B). Thus, SINE and LINE repeats have a distribution compatible with them being absent from uaRNAs and 3’ extensions in proliferating cells, but present within the pegeRNAs produced by the senescent cells.

**Figure 6:**
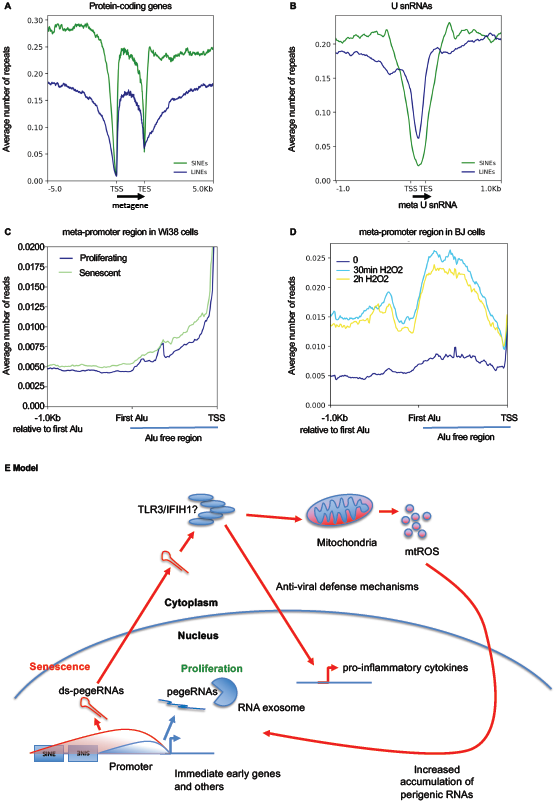
pegeRNAs from senescent cells reach into SINE- and LINE-containing regions. (A, B) Average number of either SINEs (green profiles) or LINEs (blue profiles) per 50 nucleotide bin in the neighborhood of either protein-coding genes (A) or U snRNAs (B). TSS: transcription start site. TES: transcription end site. Gene bodies are all scaled to 2Kb. U snRNA bodies are scaled to 200 nucleotides. Location of SINEs and LINEs was obtained from the Hg19 version of RepeatMasker. (C, D) At the 5260 human promoters not overlapping with coding regions of any gene described in Figure 1, the region from the TSS to the first Alu sequence was scaled to 1Kb (indicated as Alu-free region). The profile then continues 1Kb after the start of this first Alu. The average number of reads per 50 nucleotide bin (CPM normalization) was then calculated within, and upstream of this region. This was carried out for the RNA-seq data from (C) WI38 hTERT RAF1-ER human fibroblasts either proliferating or driven into senescence (Lazorthes et al., 2015) and (D) BJ cells exposed to H2O2 for the indicated times (Giannakakis et al., 2015). (E) Hypothetical model: an initial source of oxidative stress increases elongation and reduces degradation of pegeRNAs that will eventually contain sequences encoded by repeats originating from retrotransposons. Because of their abundancy, a fraction of the pegeRNAs reaches the cytoplasm (Giannakakis et al., 2015). In the cytoplasm, double-stranded RNAs generated by inverted repeats are detected by anti-viral defense mechanisms. Activation of these RNA receptors results in mitochondrial dysfunction (Djafarzadeh et al., 2011). This causes production of mitochondrial reactive oxygen species (mtROS) that maintains the hampered RNA exosome activity and feeds the inflammatory phenotype of senescent cells.

To document this, the average distribution of reads at promoters in Wi38 cells driven into senescence by oncogenic RAF initially shown Figure 1G was re-plotted to indicate the position of the first Alu upstream of the TSS (Fig. 6C). In this new plot, the region from the TSS to the first Alu was scaled to a fixed 1Kb. A similar plot was also drawn for the BJ cells exposed to H2O2 (Fig. 6D). This allowed distinguishing transcription in the Alu-free promoter region from that occurring beyond the first Alu. In both data sets, we observed that in proliferating cells, uaRNAs rarely reached beyond the first Alu, while in the senescent cells or in cells exposed to oxidative stress, transcription of regions beyond the first Alu was increased.

These observations define pegeRNAs as a source of transcripts containing transposon sequences, and we speculate that these transposons could be detected by the anti-viral defense mechanisms upregulated in senescent cells.

## Discussion

Senescent cells have many characteristic phenotypes, including growth arrest, modified chromatin structure, and secretion of proinflammatory molecules. Here, we document an additional characteristic, namely the accumulation of RNA species transcribed from regions located upstream and downstream of genes. These RNA species that we refer to as perigenic RNAs (pegeRNAs), include uaRNAs and 3’ extensions of genes. Accumulation of these RNAs in association with senescence was detected in multiple cell lines, driven into senescence via various routes, including forced expression of oncogenes, DNA damage, and replicative exhaustion.

3’ extensions of genes are removed from the main transcript during the process of mRNA polyadenylation or U snRNA maturation. The cleaved product is then degraded by the RNA exosome. Likewise, this RNA degradation machinery clears the cells of uaRNAs (Ogami, Chen et al., 2018). Accordingly, we observed reduced expression of RNA exosome subunits in two of the senescent cell lines we examined, namely the Wi38 and the IMR90 cells driven into senescence by oncogenic RAF and RAS, respectively. In both cases, expression of the DIS3L gene coding for the catalytic subunit of the cytoplasmic RNA exosome was affected. Expression of this subunit was also down-regulated in the cells exposed to chronic oxidative stress in the mouse model inactivated for Mof. In the IMR90 and in the mouse cells, expression of the regulatory subunits EXOSC3 and EXOSC9 were also down-regulated. In contrast, EXOSC4 was upregulated in all three cellular models, for reasons which are as yet unclear.

In the other senescent cells we examined, expression of DIS3L was down-regulated less than two-fold. We speculate that in these cells, the reduced RNA decay may also involve post-translational mechanisms. The existence of such mechanisms is suggested by the very rapid accumulation of pegeRNAs (30 min.) observed in cells exposed to H2O2 (this paper and original analysis by (Giannakakis et al., 2015)). Similarly, the half-life of unstable non-coding RNAs was previously shown to be drastically increased in cells exposed to oxidative stress (Tani, Numajiri et al., 2019).

To estimate RNA decay efficiency, we took advantage of the differential sensitivity of stable and unstable mRNAs to changes in the activity of RNA degradation machineries. This approach was verified in cells with reduced RNA exosome activity caused by depletion of EXOSC3. Calculating the ratio between reads mapping to unstable RNAs and stable RNAs (Lo-Hi RNA stability ratio) allowed us to address changes in RNA decay efficiency even in RNA-seq datasets of insufficient depth for the detection of the rare pegeRNAs or carried out on poly(A) selected libraries. This Lo-Hi RNA stability ratio may be of general interest as a first approach to RNA decay in, for example, cancer cells.

Increased Lo-Hi RNA stability ratio was associated with increased expression of OAS1 and OAS2 in essentially all the senescent cells we examined. We speculate that this is a manifestation of cells facing accumulation of double-stranded RNAs in the cytoplasm. While this could not be verified in senescent cells as no cytoplasmic RNA-seq data of sufficient quality was available, an earlier study has shown that pegeRNAs are extensively detected in the polysomes of cells exposed to H2O2 (Giannakakis et al., 2015). Possibly, the high stability of these RNAs in cells exposed to stress may increase the probability of their export from the nucleus to the cytoplasm.

Examination of the transcriptomes also provided indications on the receptors possibly involved in the detection of the stabilized RNAs. Indeed, we noted that upregulation of the interferon-inducible OAS1 and OAS2 was frequently associated with increased expression of TLR3, IFIH1/MDA5, and DDX58/RIG-I genes, all RNA sensors upstream of the interferon pathway. Although the regulation of TLR3, IFIH1/MDA5, and DDX58/RIG-I is poorly documented, their increased transcription may reflect an increased need for RNA detection in the lengthy process of entering senescence.

Surprisingly, the Wi38 cells driven into senescence by RAF or RAS did not activate TLR3. Instead, we noted increased expression of the gene encoding the NLRP3 inflammosome sensor protein. Unlike TLR3 that detects double stranded RNAs, NLRP3 was reported to be activated by Alu RNAs. RNAs encoded by these retransposons have previously been shown to stimulate secretion of cytokines (Hung et al., 2015). This emphasizes the possible role of repeated motifs originating from retrotransposons in the activity of pegeRNAs. In senescent cells and cells exposed to oxidative stress, pegeRNAs will harbor sequences from transposable elements as the transcripts extend beyond the repeat-free promoter and termination regions. In turn, these sequences will potentially be subject to direct detection by for example Ro60/TROVE2 that bind inverted Alu sequences (Hung et al., 2015), or will fold into double stranded RNAs if the transposons form inverted repeats.

Among the senescent cells we examined, we also identified four cases for which the Lo-Hi RNA stability ratio was either unchanged or decreased, namely the fibroblasts IMR90 and HCA-2, and the keratinocytes driven into senescence by gamma irradiation, and the IMR90 fibroblasts subject to replicative senescence. These cells did not activate OAS1 and OAS2, allowing us to conclude that activation of the OAS/RNASEL pathway is not a signature of senescence, but rather a signature of reduced RNA decay, and that this transcriptional pattern is to be found in only a subset of senescent cells.

An essential and well-described avenue to senescence is the accumulation of DNA in the cytoplasm, activating the cGAS/STING1 pathway and triggering inflammation. Accumulation of DNA in the cytoplasm was reported linked to reduced accumulation of the RNAses TREX1 and DNAse2. At the transcriptional level, we did indeed observe reduced expression of these genes in some of the cell types we examined. But in several other cases, we observed increased rather than decreased expression of DNASE2. A possible explanation for this is that the DNA accumulating in the cytoplasm needs to be processed by DNASE2 to be detected by the DNA receptor TLR9 (Chan, Onji et al., 2015). This may also explain why in some cells, we observe upregulation of TLR9.

The overall conclusion from examination of RNA-seq data from senescent cells is that some cells seem confronted just with ectopic RNA accumulation, while others seem to manage both ectopic RNA and DNA, as suggested by the parallel expression of both the TLR3 and the TLR9 genes. This leads us to propose that the inflammatory phenotype of senescent cells is triggered by multiple pathways as a function of the context. This would imply that it may be impossible to define a unique transcriptional signature of senescent inflammation other than the end-point activation of proinflammatory cytokines.

The final question raised by our observations is whether pegeRNAs could participate in the onset of inflammation. Inactivation of Exosc3 in mouse ES cells triggered several senescence associated phenotypes, including activation of Cdkn1a/p21 and Cdkn2a/p16, activation of the p53 pathway, and reduced expression of multiple markers of cell cycle progression, all in favor of a triggering effect of pegeRNA. In addition, inactivation of Exosc3 in these cells resulted in reduced expression of mitochondrial genes. Interestingly, medical data recapitulate this observation. Indeed, patients suffering from pontocerebellar hypoplasia, a disease affecting the development of the brain and due to a mutation in the *EXOSC3* gene were also described as showing signs of mitochondrial dysfunction (Schottmann et al., 2017). In parallel, patients with mutations in *EXOSC2* are subject to premature ageing (32). TLR3, IFIH1/MDA5, DDX58/RIG-I, and NLRP3 are all connected with mitochondria (Djafarzadeh, Vuda et al., 2011, Zhou, Yazdi et al., 2011), while mitochondrial dysfunction is a source of oxidative stress, that in turn is a trigger of pegeRNA accumulation. We suggest that together, these elements compose a possible feedback loop in which accumulation of pegeRNAs triggers anti-viral defense mechanisms causing production of mitochondrial reactive oxygen species that in turn nurtures both the inflammatory response and the reduced RNA turnover. Once initiated, such a process could be an irreversibly driver of cellular senescence (Model Figure 6A).

## Acknowledgements

We thank M. Ricchetti for critical reading of the manuscript. The work was supported by grants from Institut Pasteur, LABEX REVIVE 10-LBX-73, and Agence Nationale de la Recherche 16-CE-15. C. Mann was supported by ANR 17-008-02.

## Material and methods

### Tissue culture

WI38 hTERT RAF1-ER cells, which are immortalized by hTERT expression and contain an inducible RAF1 oncogene fused to an estrogen receptor (ER), were maintained in minimum essential medium (MEM) supplemented with gluta-mine, non-essential amino acids, sodium pyruvate, penicillin-streptomycin, and 10% fetal bovine serum in normoxic culture conditions (5% O2) (Jeanblanc, Ragu et al., 2012). For the induction of oncogene-induced senescence, cells were treated with 20nM 4-HT (H7904, Sigma) for 3 days.

### Western blotting

Cells were lysed with lysis buffer (50 mM Tris-HCl pH 7.5, 150 mM NaCl, 1% Triton X-100, 0.1% SDS, 1 mM EDTA, and protease/phosphatase inhibitor mixture (Roche)) and sonicated (Bioruptor, Diagenode). The protein concentration was determined by Bradford assay, and 20 µg of protein were boiled in the presence of NuPAGE LDS sample buffer (Invitrogen NP0007) and NuPAGE Sample Reducing Agent (Invitrogen NU0004) at 95 °C for 5 min, resolved by SDS-PAGE (4-12% Criterion(tm) XT Bis-Tris Protein Gel, Bio-Rad), and transferred to nitrocellulose membrane (Bio-Rad). Staining with ATX Ponceau S Red (Sigma-Aldrich, 09189) was used as a further marker of protein content. The membrane was then blocked with 5% non-fat milk in phosphate buffer saline (PBS)-0.1% Tween20 (Sigma-Aldrich, P1379) for 1h at RT and probed with specific primary antibodies (α-p21 (1:500, BD Biosciences, 556430), DIS3L (1:500, ab89042) and actin (1:1000, Sigma-Aldrich, A2103) overnight at 4 °C. After three washes in PBS containing 0.1% Tween20, the membrane was incubated with anti-Rabbit or anti-Mouse IgG HRP secondary antibodies for 1h at RT and revealed by chemiluminescence respectively. Detection was performed using Chemidoc(tm) MP imaging system (Bio-Rad). Experiments were done in triplicate, and a representative Western blot was shown. Western blot bands quantification were done using the ImageJ software.

### Data download

RNA-seq raw fastq files were downloaded from the Gene Expression Omnibus (GEO) on the NCBI database with the SRA toolkit (http://ncbi.github.io/sra-tools/). The files were retrieved from GSE55172, GSE77784, GSE81662, GSE85085, GSE100535, GSE108278 and GSE130727. The fastq files from the analysis E-MTAB-5403 were downloaded from ArrayExpress on the EBI database.

### Mapping

For the mapping, SHRiMP were used (v2.2.3) (David, Dzamba et al., 2011) for the dataset GSE55172 (colorspace reads) (parameters: -o 1 --max-alignments 10 --strata) while for the others, mapping were done with STAR (v2.6.0b) (Dobin, Davis et al., 2013) (parameters: --outFilterMismatchNmax 1 --outSAMmultNmax 1 -- outMultimapperOrder Random --outFilterMultimapNmax 30). Mapping were done against the reference human genome (hg19 homo sapiens primary assembly from Ensembl) for the EBI dataset, GSE55172, GSE81662, GSE85085, GSE108278, GSE130727 and against the reference mouse genome (mm9 mus musculus primary assembly from Ensembl) for the GSE77784 and GSE100535). The SAM files were then converted to the BAM format and sorted by coordinate with samtools (v1.7) (Li, Handsaker et al., 2009).

### Data observations

Bigwigs files were generated with bamCoverage (parameter: --normalizeUsing CPM) from Deeptools (v3.1.3) (Ramirez, Ryan et al., 2016). All observations were done using the Integrative Genomics Viewer software (IGV) (Robinson, Thorvaldsdóttir et al., 2010).

### Differential gene expression

The package Subread (v1.28.1) (Liao, Smyth et al., 2014) for R (v3.4.3) were used to count the uniquely mapped reads based on a gtf annotation file for hg19 or mm9 from Ensembl. Then the package DESeq2 (v1.18.1) (Love, Huber et al., 2014) were used to make the differential gene expression analysis and principal component analysis (PCA). P-values from the differential gene expression test were adjusted for multiple testing according to the Benjamini and Hochberg procedure. Only genes with an adjusted p-value lower than 0.05 were considered differentially expressed. GO term analysis were performed on these differentially expressed genes with Enrichr (Kuleshov, Jones et al., 2016).

### Heatmaps, and profiles

Reads inside upstream gene regions and downstream U snRNA regions were quantified using featureCounts (v1.6.1) from the Subread suite (Liao et al., 2014). Then, from these counts, matrices were generated with the tool computeMatrix reference-point (parameter: --referencePoint TSS for the observations of the regions upstream of the genes or --referencePoint TES for the observations of the downstream regions of U snRNA). Profiles were obtained using plotProfile (parameter: --perGroup) and heatmaps using plotHeatmap (default paramaters) from the Deeptools suite (Ramirez et al., 2016).

### Calculation of the Lo-Hi RNA stability index

The package Subread (v1.28.1) (Liao et al., 2014) for R (v3.4.3) were used to count the uniquely mapped reads based on a list of either highly unstable (t1/2<2h) or high stable (t1/2>10h) mRNAs from (Tani et al., 2012). For each feature, the count was normalized using the TPM method. Then, for both condition, the sum of every TPM has been done and the highly unstable mRNAs was divided by the high stable mRNAs, giving us a value for this Lo-Hi RNA stability index. Bed files of the highly unstable and high stable mRNAs are available as supplemental data.

## Figure legends

**Sup. Figure 1:**
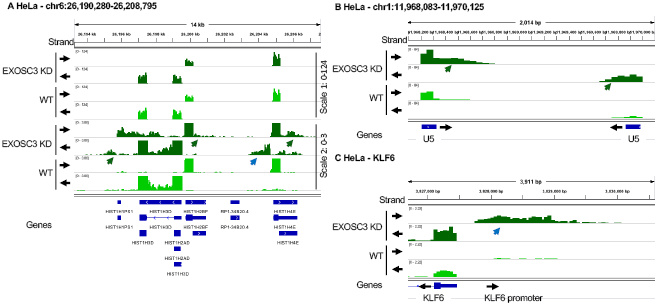
Senescent WI38 cells accumulate pegeRNAs. (A-C) RNA-seq data from HeLa cells transfected either with siLuc (control) or siEXOSC3 (Schlackow, Nojima et al., 2017). RNA-seq reads were quantified, normalized by the CPM method and displayed with IGV. Black arrows indicate the orientation of the transcription. Green arrows indicate 3’ extensions, blue arrows, uaRNAs.

Sup Table 1A: List of genes accumulating uaRNAs 2-fold more in senescent as compared to proliferating Wi38 cells.

Sup Table 1B: List of snRNA genes accumulating downstream reads 2-fold more in senescent as compared to proliferating Wi38 cells.

**Sup. Figure 2:**
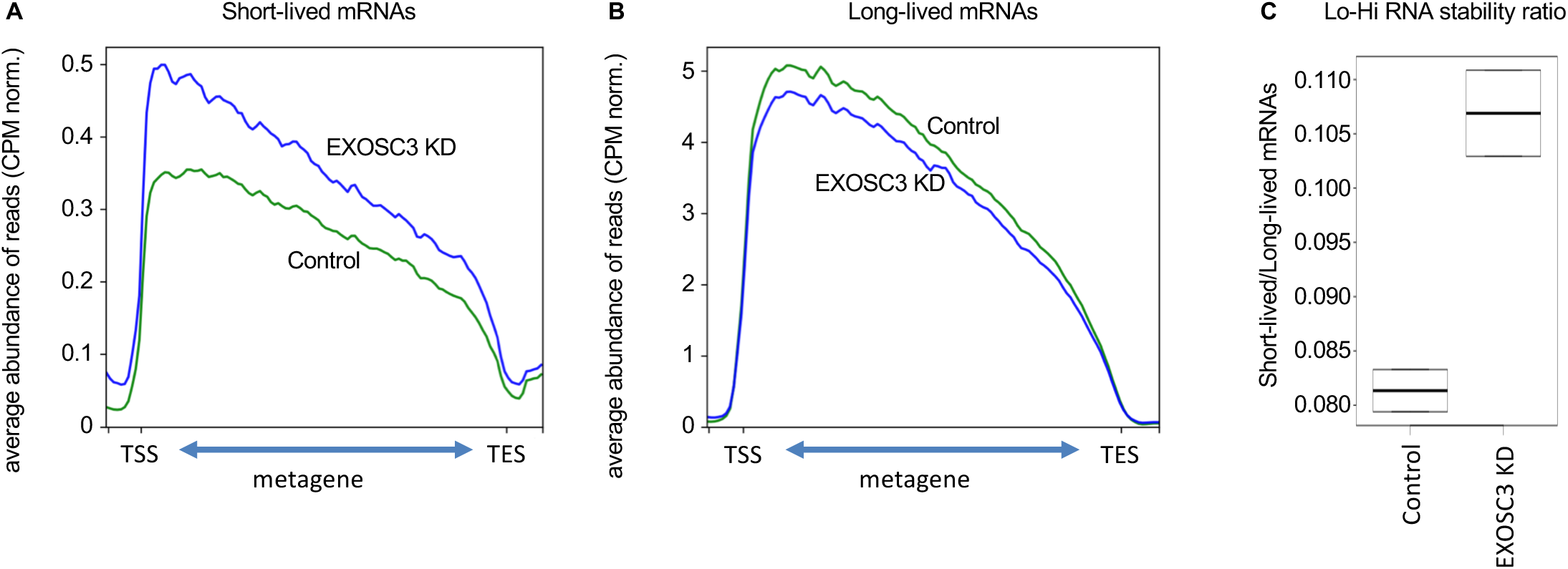
Reduced expression of RNA exosome subunits in senescence. RNA-seq data from HeLa cells transfected either with siLuc (control) or siEXOSC3 (Schlackow et al., 2017). (A, B) Average profile of reads mapping to short-lived (less than 2h) or long-lived (more than 10h) mRNAs as listed in (Tani et al., 2012), either in Control (green) or in EXOSC3 knock-down (blue) cells as indicated. Read counts were normalized by CPM. (G) Ratio of the number of reads mapping to short-lived (less than 2h) over the number of reads mapping to long-lived (more than 10h) mRNAs.

**Sup. Figure 3:**
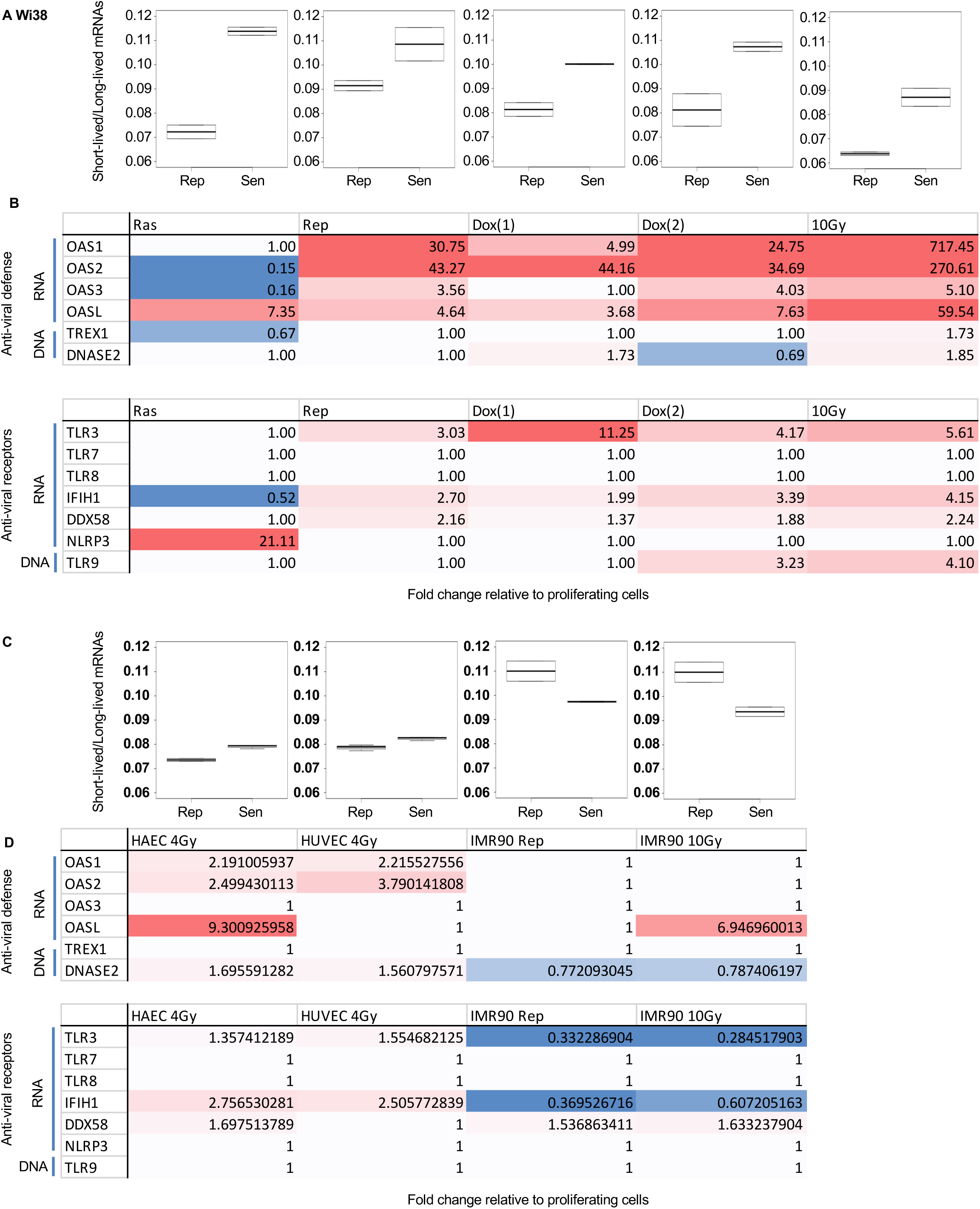
Reduced RNA decay in senescent cells of various origin RNA-seq data from (Casella et al., 2019), n=2 for all experiments. (A) Wi38 fibroblasts were driven into senescence either by forced expression of oncogenic RAS, continuous replication, or DNA damage with doxorubicin (2 experiments) or gamma irradiation (10Gy). Lo-Hi RNA stability ratio for each experiment was calculated as in Figure 2. (B) Fold change relative to proliferating cells of the indicated genes was estimated with DESeq2. Red gradient indicates upregulation (max red at 10-fold), blue indicates downregulation (max blue at 0.5). Variations with pVal>0.05 were set to 1. (C-D) HAEC and HUVEC cell lines were driven into senescence by gamma irradiation, and IMR90 cells were rendered senescent either by replication exhaustion or gamma irradiation (same controls for the two routes). (C) Lo-Hi RNA stability ratio for each experiment was calculated as in Figure 2. (D) Fold change relative to proliferating cells of the indicated genes represented as in (B).

**Sup. Figure 4:**
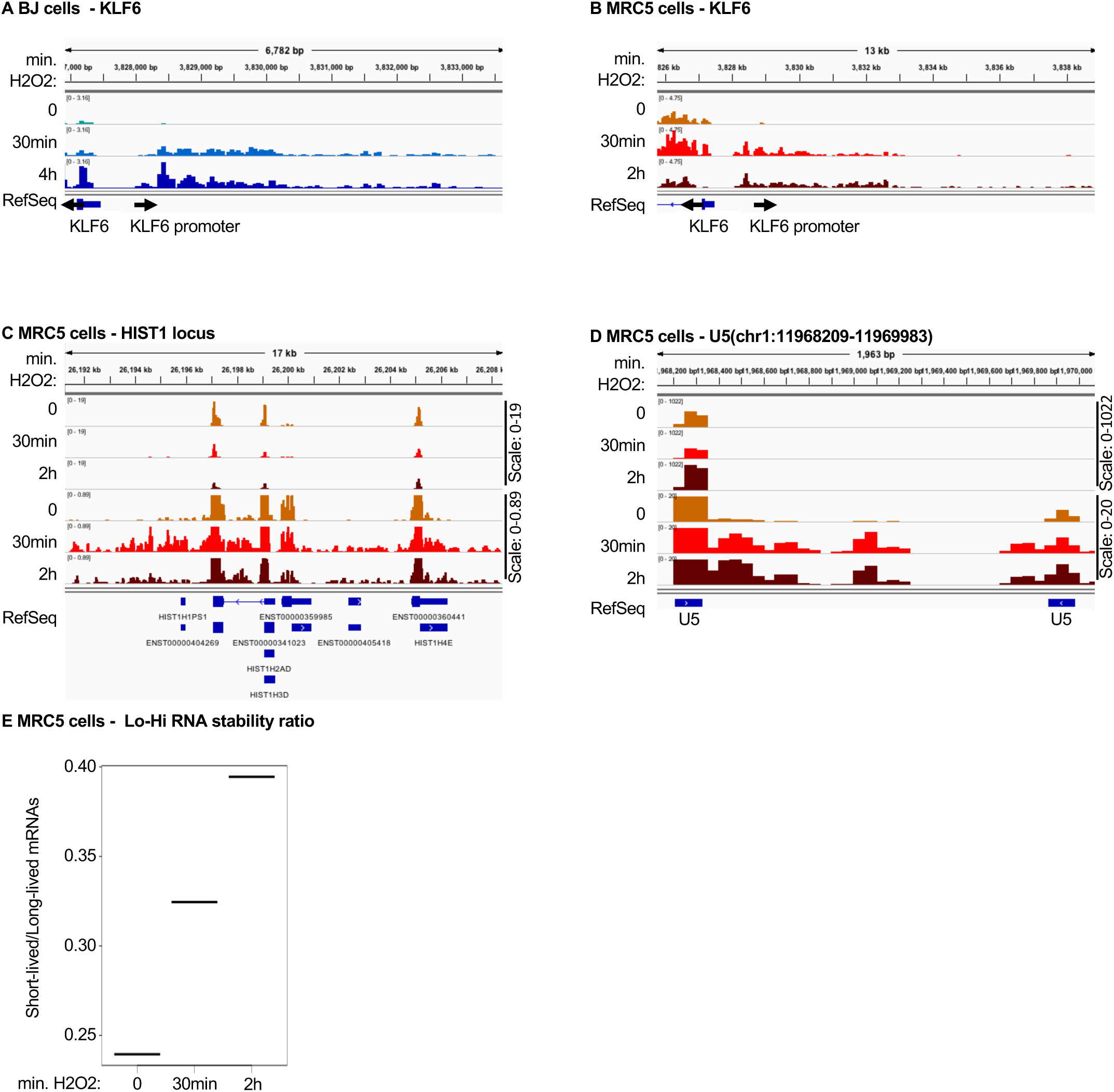
Accumulation of pegeRNAs in cells exposed to oxidative stress BJ or MRC5 human fibroblasts having been exposed to 0.2mM H2O2 for the indicated times (one sample per time point) (Giannakakis et al., 2015). (A-D) RNA-seq reads were quantified, normalized by the CPM method and displayed with IGV genome browser. (E) Lo-Hi RNA stability ratio for MRC5 cells at the indicated time points, calculated as indicated in Figure 2.

Sup. Table 4: List of snRNA genes accumulating downstream reads 2-fold more in H2O2-treated as compared to untreated BJ cells.

Sup. Table 5: differentially regulated genes associated with the GO terms in Figure 5 panel C, E, and F.

## Notes

### Competing Interest Statement

The authors have declared no competing interest.

### Summary of Updates

This version of the paper provides additional evidence for reduced RNA decay associated with cellular senescence, and explores data from a larger series of cellular models. These additional data have revealed that it is only a subset of senescent cells that display the reduced RNA decay, along with activation of anti-RNA defense mechanisms. This observation further suggests that it may be possible to make a distinction between DNA- and RNA-driven inflammatory phenotypes in the context of cellular senescence.

